# Sequence dependent co-phase separation of RNA-protein mixtures elucidated using molecular simulations

**DOI:** 10.1101/2020.07.07.192047

**Authors:** Roshan Mammen Regy, Gregory L. Dignon, Wenwei Zheng, Young Chan Kim, Jeetain Mittal

## Abstract

Ribonucleoprotein (RNP) granules are membraneless organelles (MLOs) which majorly consist of RNA and RNA-binding proteins and are formed via liquid-liquid phase separation (LLPS). Experimental studies investigating the drivers of LLPS have shown that intrinsically disordered proteins (IDPs) and nucleic acids like RNA play a key role in modulating protein phase separation. There is currently a dearth of modelling techniques which allow one to delve deeper into how RNA plays its role as a modulator/promoter of LLPS in cells using computational methods. Here we present a coarse-grained RNA model developed to fill this gap, which together with our recently developed HPS model for protein LLPS, allows us to capture the factors driving RNA-protein co-phase separation. We explore the capabilities of the modelling framework with the LAF-1 RGG/RNA system which has been well studied in experiments and also with the HPS model previously. Further taking advantage of the fact that the HPS model maintains sequence specificity we explore the role of charge patterning on controlling RNA incorporation into condensates. With increased charge patterning we observe formation of structured or patterned condensates which suggests the possible roles of RNA in not only shifting the phase boundaries but also introducing microscopic organization in MLOs.

## INTRODUCTION

Membraneless organelles (MLOs) are compartments formed in the cell which consist of a concentrated set of biomolecules without an enclosing membrane separating them from the surrounding cytosol (1–3). Many of these MLOs are assemblies consisting of RNA-binding proteins and RNA are commonly referred to as Ribonucleoprotein (RNP) granules (4). Prominent RNP granules include P bodies, stress granules, germ cell P granules, and neuronal granules which perform diverse functions that are essential for the survival of the cell. P bodies are mRNA and protein containing cytoplasmic processing bodies associated with RNA metabolism (5). Stress granules are formed when cells respond to stress by selectively including specific mRNA transcripts into granules and regulating/arresting translation (6). Germ cell P granules form during germ cell development (7), whereas neuronal granules transport mRNAs in response to specific exogenous stimuli (8). Despite the highly diverse functions, these RNP granules share a common process through which they localize their constituent proteins and nucleic acids, that is, through liquid-liquid phase separation (LLPS). Therefore, investigating the molecular mechanism underlying LLPS of biomolecules is essential to understanding how RNP granules store, process and control the activities of their constituents.

It is now well acknowledged that these MLOs are formed when a homogeneous mixture of biomolecules undergoes the physical process of LLPS to form coexisting condensed and dilute phases stabilized by a balance between entropic and enthalpic interaction (9, 10). These biomolecular condensates may appear as liquid-like droplets and allow a rapid exchange of components with the environment in a dynamic manner (11). Investigations into the drivers of protein LLPS have also shown that the disordered domains of certain proteins such as the DEAD-box helicase LAF-1 and the RNA-binding protein Fused in Sarcoma (FUS) are able to create phase separated condensates *in vitro* (11, 12), thus suggesting the importance of intrinsically disordered regions (IDRs) in facilitating the formation of RNP granules (13). Incorporation of RNA into condensed phases has been further shown to perturb the physical and chemical properties of these condensates (11, 12, 14–18). For instance, incorporating RNA into the condensed phase of the DEAD-box helicase LAF-1 protein, which is a major component of P bodies, can shift the thermodynamic phase diagram for the LAF-1 protein and its disordered RGG rich domain and change both the saturation concentration and LAF-1 diffusion inside the liquid droplets in *in vitro* experiments (11). In another experimental study varying the concentration of RNA in a co-phase separating mixture of RNA and FUS shows re-entrant protein phase separation behavior where RNA acts as a facilitator of FUS phase separation at low concentration but on increasing the RNA concentration beyond a certain point we see RNA acting as a disruptor of protein LLPS (12, 19). These studies have brought forth how RNA plays an essential role as a modulator of protein phase separation and condensate properties once it is incorporated into the protein-rich phase.

Despite these advances in establishing the role RNA plays in the formation and function of RNP granules, there currently exists a lack of understanding about the fundamental molecular driving forces driving RNA-protein co-phase separation and its more nuanced aspects like the microstructural organization of different components within the condensates formed. Although remarkable progress has been recently made in providing insights into the role of a protein’s sequence in controlling its LLPS (20–22), how the protein sequence plays a role in modulating LLPS of protein-RNA mixtures and the material properties of the condensates formed is not well established (19, 23, 24). Part of this stems from the lack of techniques that can provide in-depth information about the RNA-protein interactions driving RNP granule formation and high spatiotemporal resolution on the molecular organization of protein and RNA within the condensate (12, 25, 26). One can attempt to obtain such in-depth information from *in silico* atomic resolution simulation techniques (20, 27, 28) but studying a macroscopic phenomenon like phase separation would require considerable computational resources making this method quite expensive and prohibitive. This prompts us to look into computational approaches based on coarse-grained (CG) models which can allow investigations into the formation of biomolecular condensates and to provide molecular-level details necessary to develop theories of phase separation making CG models an integral part of the biophysical toolkit to study phase separation (29–32). We have previously developed a CG modeling framework based on the amino acid hydropathy (HPS model) to study sequence determinants of protein phase separation that does not require input from experimental data (33, 34). The HPS model is based on defining nonbonded interactions between amino acid pairs using the hydropathy values of the naturally occurring twenty amino acids (33, 35). This model was recently extended to post-translationally modified amino acids as well (36).

In the present work, we first define parameters for RNA nucleotides within the HPS modeling framework using a one bead per nucleotide CG representation of RNA. We then test the new model to study the co-phase separation of the N-terminal disordered RGG domain of the LAF-1 protein (hereafter referred to as LAF-1 RGG) and RNA molecules with the most recent experimental data available (11). The model provides reasonable agreement with the experiment regarding the effects of RNA on modulating LAF-1 RGG phase separation. We then utilize this new CG model to study the co-phase separation LAF-1 RGG protein variants with the same sequence composition but different arrangement of charged amino acids used in our previous work (37). Consistent with the expectations from previous work (31, 37), we find that the phase behavior can be significantly perturbed by changes in the protein sequence, and this change in behavior also alters co-phase separation. Most importantly, we observe a significant change in terms of the co-localization of protein and RNA molecules within the condensate. For the charge segregated variants of the LAF-1 RGG protein, RNA adsorbs on the protein condensate surface rather than mixing evenly throughout as in condensates of WT LAF-1 RGG and RNA mixtures. We provide interpretation for the observed behavior based on the potential of mean force between pairs of protein and RNA molecules. Our work here attempts to provide evidence linking competing intermolecular interactions with the sub-compartmentalization inside MLOs allowing us to better understand what determines the morphology of MLOs.

## MATERIAL AND METHODS

### HPS model for proteins

Our CG modeling approach for proteins uses a one bead per amino acid level of resolution, and the twenty naturally occurring amino acids are identified in the model by the following characteristics: the mass, charge, diameter (*σ*) (38), and hydropathy (*λ*) of the amino acid. The *λ* value is derived from the partial charges of atoms belonging to an amino acid in an all-atom force field and scaled to range from 0 (Arginine) to 1 (Phenylalanine), which is based on the approach proposed by Kapcha and Rossky (35). In the CG energy function, we have three types of interactions: bonded, electrostatic, and short-range pairwise interactions. The bonds between consecutive amino acids in the protein sequence are modeled using a harmonic potential with a spring constant of 10 kJ/Å^2^ and a bond length of 3.8 Å. The electrostatic interactions are modeled using the Debye-Hückel (DH) electrostatic screening term (39),

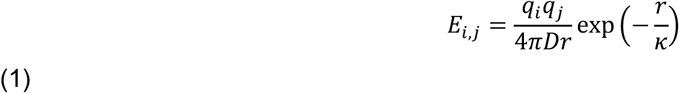

in which к is the Debye screening length, and *D* = 80 is the dielectric constant of the solvent. We set к = 10 Å, corresponding to solution conditions at 100 mM salt. The short-range pairwise interactions are modeled by the Ashbaugh-Hatch functional form (40) as

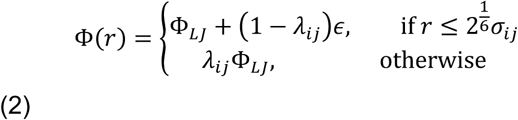

where Φ_*LJ*_ is the standard 12-6 Lennard-Jones potential given by

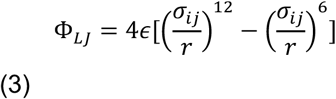

Both the hydropathy (*λ*_ij_) and the diameter (*σ*_ij_) are set to be the arithmetic average of the values for the interacting pair of amino acids (*i,j*). A value of *λ*_ij_ close to 1 represents strong attractive interactions whereas a value of 0 represents weak repulsive interactions between the two amino acids. The interaction parameter (*ε*) in eq. 3 is a free parameter and was adjusted to reproduce the experimentally measured radius of gyration of IDPs as 0.2 kcal/mol (33).

### HPS model for RNA

Our purpose for building a new RNA model, as opposed to using an existing model (41–43), is to capture the qualitative features of the RNA-protein co-phase separation behavior with the simplest possible CG representation that is also consistent with our protein model. Hence, we decided to model RNA with each nucleotide represented as a single particle (Fig. 1A). The potential energy also consists of the bonded, electrostatic and short-range pairwise interaction terms, the same as that of the disordered protein but with different parameters (Table 1). A spring constant of 10 kJ/Å^2^ and a bond length of 5 Å is used for the bonded term, in which the bond length is derived using the average backbone-backbone distance for single- and double-stranded DNA (44).

**Table 1.** RNA Model parameters for CG beads, each representing a single nucleotide.

| Nucleotide | Mass (amu) | Charge (1) | Diameter, $\sigma$ (Å) | Hydrophathy, $\lambda$ (1, 35) |
| --- | --- | --- | --- | --- |
| Adenosine | 329.2 | -1.0 | 8.44 | -0.054 |
| Cytidine | 305.2 | -1.0 | 8.22 | -0.027 |
| Guanosine | 345.2 | -1.0 | 8.51 | -0.189 |
| Uridine | 306.2 | -1.0 | 8.17 | -0.027 |

**Figure 1:**
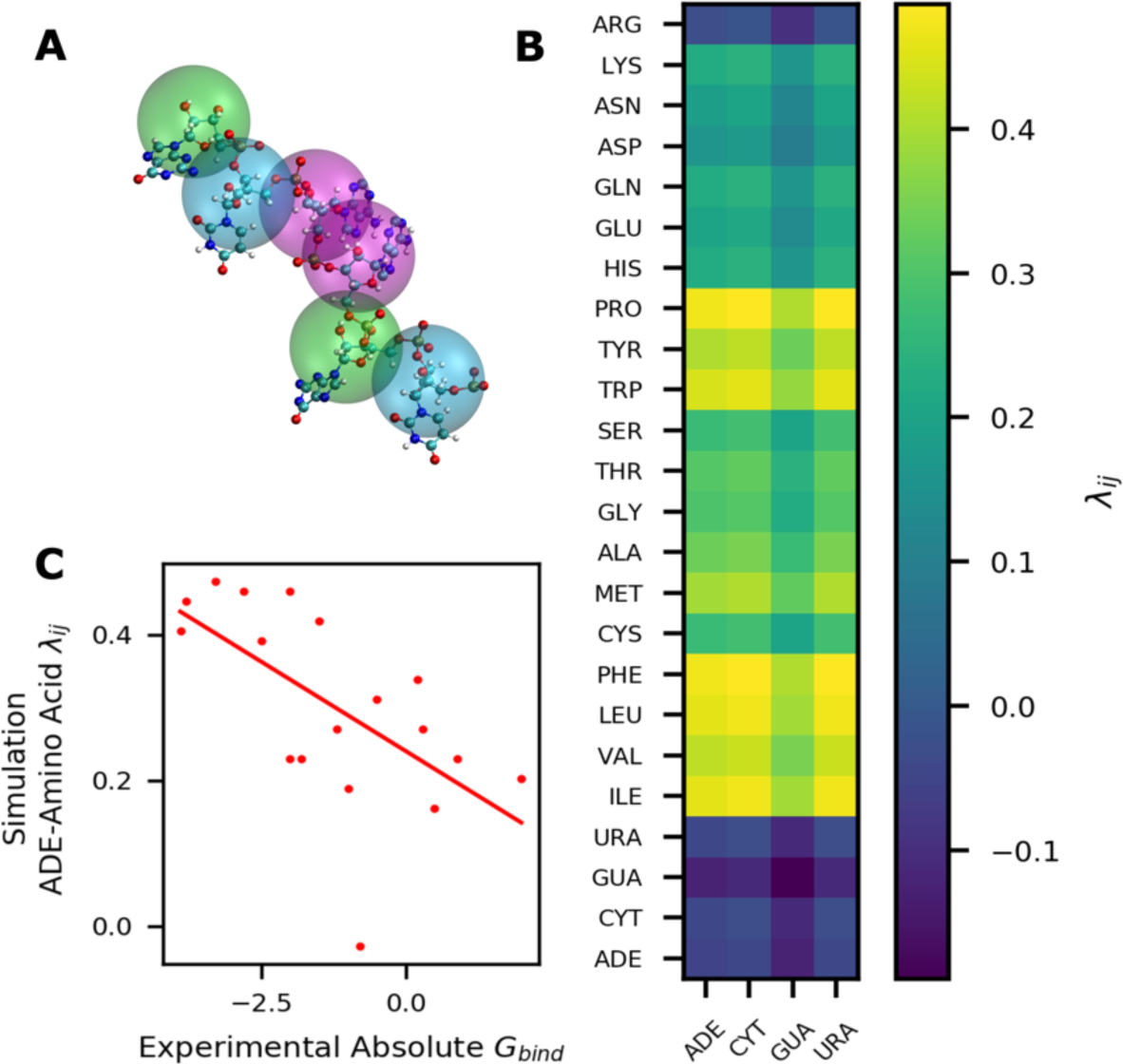
RNA Coarse Grained model details and comparison: (A) One bead per nucleotide CG representation of RNA; (B) Average hydropathy values for pairs of twenty amino acids and four RNA base pairs in the model; (C) Comparison of RNA-protein hydrophobic interaction strengths between the proposed RNA model and existing absolute binding free energy (kJ/mol) data from all-atom umbrella sampling simulations (46).

The primary mode of interactions between RNA and proteins is expected to be via electrostatics (24) due to the negatively charged backbone which is captured in our model by placing a negative charge on each of the nucleotide beads. In addition, nucleotides can interact with all other components via non-electrostatic interactions that are captured by the short-range potential (eq. 2). The hydropathy value of each nucleotide is assigned in a similar fashion as the amino acids using the atomic scale partial charges from the OPLS all-atom RNA force field (45). The RNA nucleotide hydropathy values on a normalized scale (based on amino acids) are found to be negative (*λ* < 0), which reflects repulsive short-range interactions. The arithmetic average between a pair of CG particles (i.e. twenty amino acids or four RNA nucleotides) is used for characterizing the short-range pairwise interactions. As shown in Fig. 1B, most of the protein-RNA short range interactions are weakly attractive with the most significant attraction between hydrophobic amino acids and nucleotides. Our hydrophobic interaction strengths were in acceptable agreement with reported absolute binding free energy data between nucleotides and amino acids (Fig. 1C) (46).

### Simulation strategy for sampling protein-RNA multicomponent phase separation

Even with the current advances in simulation methodology and the CG nature of the models being used, it is nearly impossible to apply standard free-energy based techniques to sample the phase behavior of long-chain off-lattice polymers (20, 28–31). We and others have been using co-existence simulation methodology (51) to sample the phase behavior of proteins undergoing LLPS successfully and efficiently (20, 32). Here, we use the same strategy, as shown in Fig. 2A, wherein a cuboid shaped periodic box or a ‘slab’ is simulated with the shorter x and y dimensions of equal length while the z dimension is extended to create a low-density phase (Fig. 2A). The use of planar interfaces in a slab configuration as opposed to a spherical droplet geometry is effective in reducing the system-size effects in calculating the densities of co-existing phases (30, 50, 51). By conducting such simulations for many temperatures, we can use the co-existing densities (Fig. 2B) to map the binodal (limits of thermodynamic stability) and the critical temperature as well as density values for the phase separating component (Fig. 2C). As in previous work, we conduct an initial equilibration of our system in a cubic simulation box consisting of all components in the NPT ensemble to reach a box size of about 15 nm in each dimension. We then extend the box z-dimension to 280 nm, and conduct coexistence simulations in the NVT ensemble for 5 μs. The temperature is controlled using the Langevin thermostat in the low friction limit. All the simulations are conducted with the HOOMD-Blue package to take advantage of its capabilities to speed-up calculations using the graphics processing units (GPUs) (52). For more details about the methodology and the analysis, we refer the readers to our previous work (33, 53).

**Figure 2:**
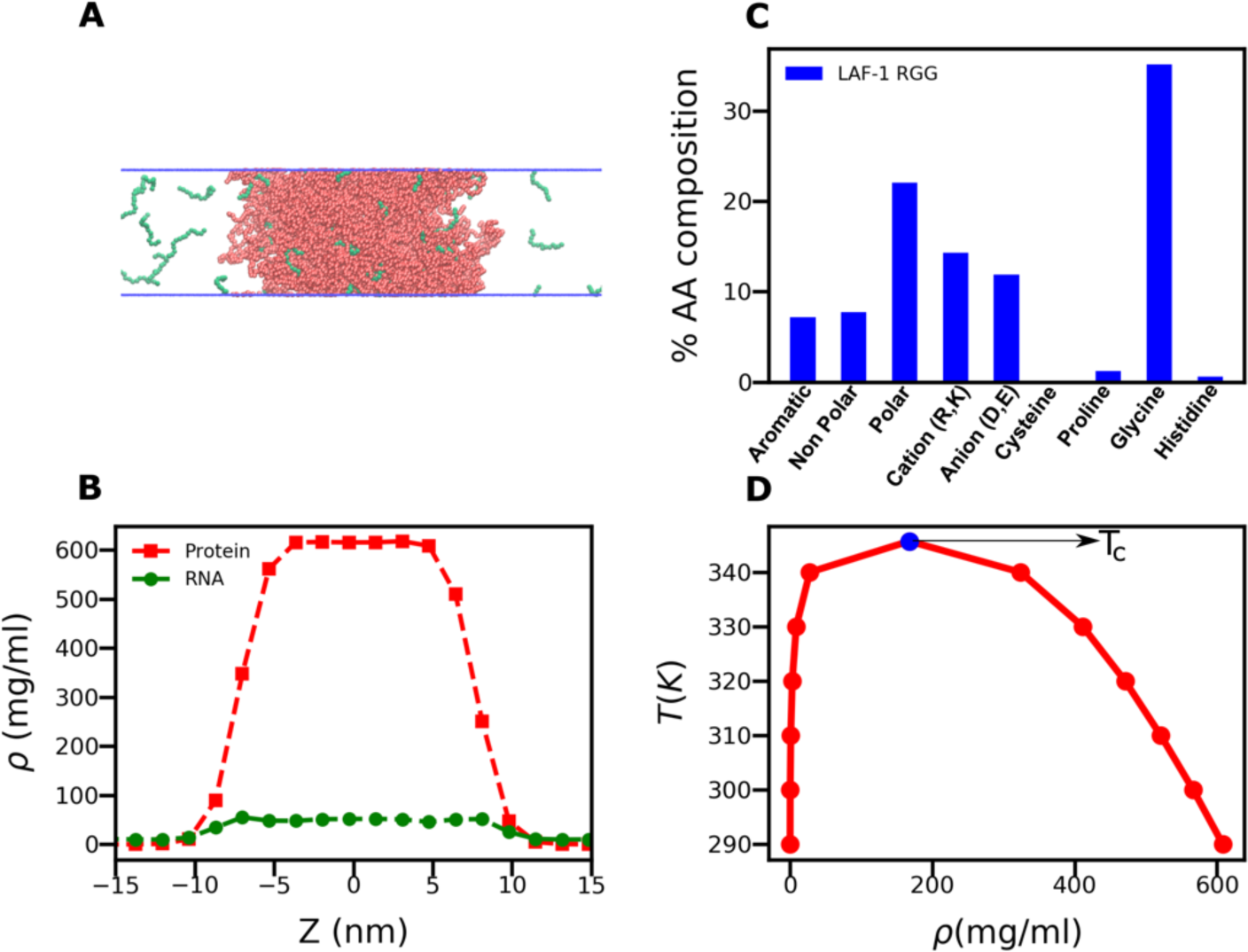
Slab simulation methodology: (A) Slab configuration for CG simulations; (B) Density profiles of LAF-1 RGG and RNA sampled over the course of a slab simulation centered on the largest cluster formed; (C) Amino acid composition of the LAF-1 RGG WT sequence. (D) Binodal phase diagram showing temperature regime over which phase separation is possible.

## RESULTS AND DISCUSSION

### RNA perturbs LAF-1 RGG phase separation in a concentration-dependent manner

We conducted coexistence simulations of the disordered LAF-1 RGG protein sequence with poly-Adenosine (poly-A)RNA of length 15 nucleotides (A_15_) at a range of temperatures and concentrations of RNA,keeping the total number of LAF-1 RGG molecules fixed. In our coexistence simulations, LAF- 1 RGG forms a protein-rich condensed phase in equilibrium with the dilute vapor-like phase (54), whereas RNA alone is not able to phase separate, even at very high concentrations (∼36mg/ml) due to electrostatic repulsion. However, in the case of a binary mixture of LAF-1 RGG and A_15_, RNA molecules are preferentially found inside the LAF-1 RGG-rich phase rather than the dilute region of the simulation box (Fig. 2B). We therefore quantify the preference of RNA molecules by calculating the partition coefficient of the RNA molecules - the ratio of the concentration of a component in the dense phase by that in the dilute phase. A high partition coefficient suggests the RNA molecule prefers to co-phase separate with the proteins whereas a small value suggests the RNA molecule tends to stay dispersed in the protein-deficient phase. The partition coefficient of the A_15_ RNA can reach values of up to ∼50. This is likely due to the presence of a relatively large number of cationic Arginine residues in the LAF-1 RGG sequence (Fig. 2D) which can form favorable interactions with the anionic RNA beads.

To observe the effect of RNA on the thermodynamics of LAF-1 RGG phase separation, we calculated the coexistence densities of protein at different temperatures and RNA solution concentrations (Fig. 3A). With increasing RNA concentration, the right arm of the binodal that represents protein density in the condensed phase shifts to the left. This suggests that the protein concentration inside this dense phase decreases which is presumably due to the increasing RNA concentration inside. This reduction in protein concentration with increasing RNA in the case of LAF-1 RGG is also consistent with previous experimental work on the same system by Elbaum-Garfinkle *et al* (23). We note that the total biomolecular concentration (protein + RNA) in the condensed phase is mostly unchanged in our simulations at a set temperature.

**Figure 3:**
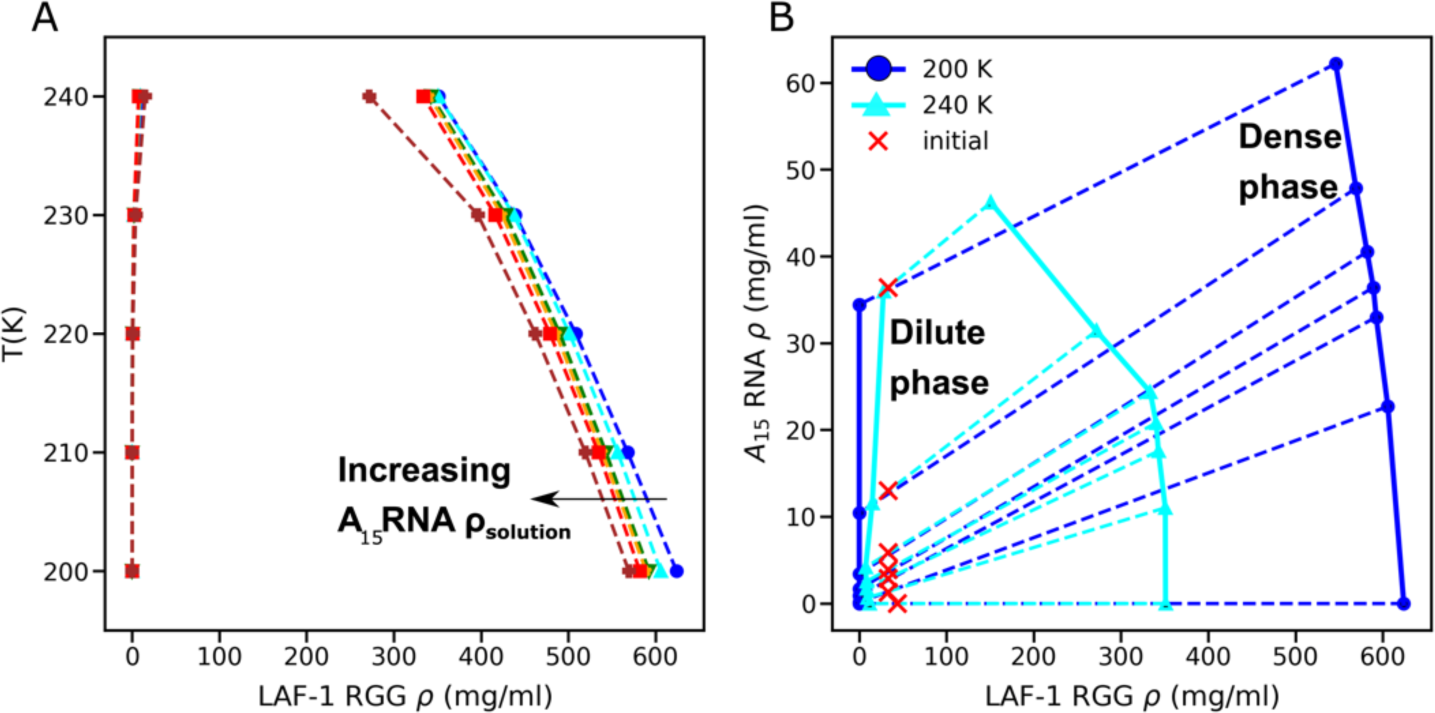
RNA and LAF-1 RGG co-phase separate to form RNA modulated protein condensates: (A) LAF-1 RGG binodal phase diagram shifts with increasing RNA solution concentration. (B) Binary phase diagram from simulations with varying solution RNA concentrations (colored (dashed) tie lines) joining the coexistence concentrations of RNA and protein at different temperatures (colored (solid lines) phase envelopes). Red X’s indicate total concentrations of simulation systems. Tie lines will always pass through these as long as the system separates into two phases.

Next, we observe the degree to which RNA is incorporated into the condensates by using the A_15_ RNA and protein concentrations from both the high- and low-density coexisting phases to construct the binary thermodynamic phase diagram (Fig. 3B). The region enclosed by these two lines is the two-phase coexistence region. If the state point as a function of LAF-1 RGG and A_15_ concentrations falls within the coexistence region, the system will phase separate into two phases with relative concentrations of protein and A_15_ RNA dictated by the tie lines joining this point to both dense and dilute phase arms of the multicomponent phase diagram (55). Based on the computed phase diagram in Fig. 3B, we observe a co-phase separation behavior which is referred to as scaffold-client type in the recent LLPS literature (10, 31), in which the scaffold can undergo phase separation on its own and recruit client molecules (which cannot phase separate alone) into the dense phase. At the given conditions, the “scaffold”, LAF-1 RGG can phase separate on its own into a protein-rich phase due to sufficiently strong homotypic attraction. This then recruits the “client” A_15_ RNA molecules for which homotypic interactions are either too weak or repulsive, but the heterotypic interactions between the scaffold and client molecules are sufficiently attractive (10, 56). The qualitative shape of the binary phase diagram remains unchanged with increasing temperature and reflects a shrinking coexisting region when approaching the critical temperature for LAF-1 RGG with an upper critical solution temperature (UCST) (34).

So far, we have mostly focused on the condensed phase arm of the phase diagram. From a biological standpoint, it is quite important to quantify the changes in the scaffold (LAF-1 RGG) concentration of the low-density phase, also referred to as the saturation concentration (C_sat_), with the changes in client (RNA) solution concentration. The significance of C_sat_ is that this is the minimum protein concentration required to undergo LLPS, so it is important to understand how the presence of RNA can be used to modulate this. In Fig. 4, we show how C_sat_ changes with increasing RNA concentration, At low RNA concentrations, there is an initial decrease in the C_sat_ values (by as much as a factor of two) highlighting the LAF-1 RGG’s enhanced propensity to phase separate in the presence of RNA. Similar LLPS behavior has also been observed previously in many other cases (19, 57, 58) which in this case is due to favorable attraction between cationic Arginine residues and anionic nucleotides.

**Figure 4:**
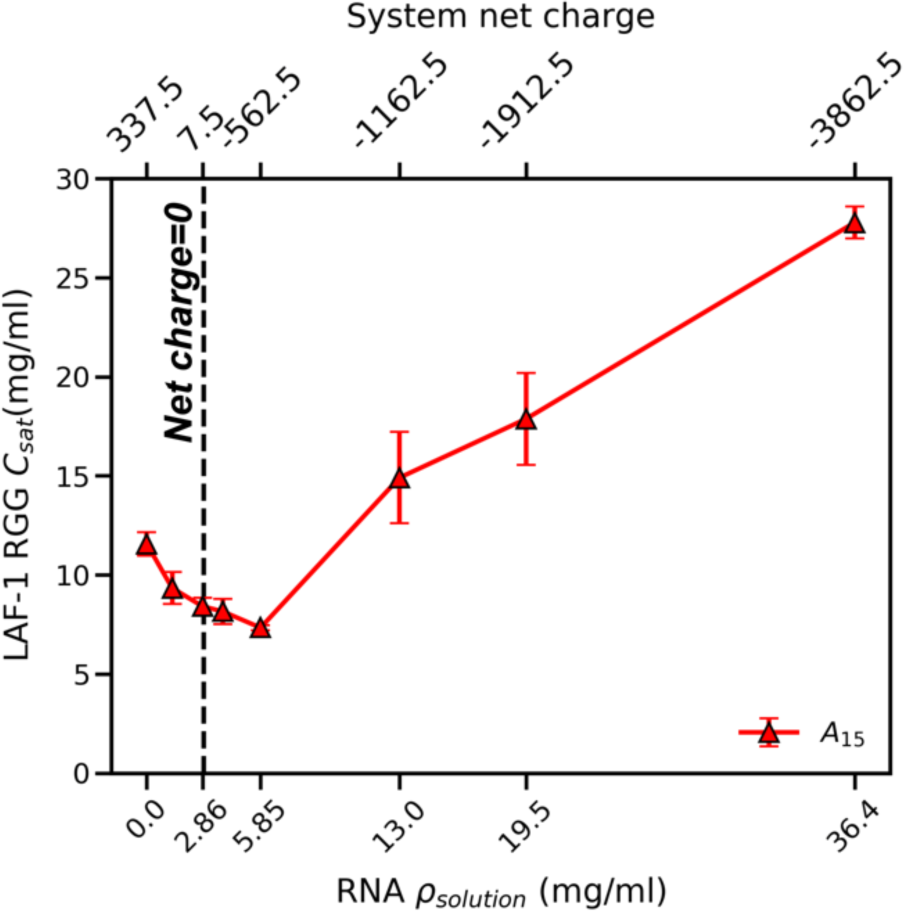
RNA shows concentration dependent protein LLPS promoter/disruptor roles for different chain lengths: Effect of increasing RNA solution concentration on LAF-1 RGG saturation concentration shows switching between RNA LLPS promoter/disruptor roles (non-monotonic curve). Vertical line represents a net charge = 0 system.

Upon a further increase in the RNA concentration beyond a certain threshold value, we observe an increase in C_sat_, highlighting LAF-1 RGG’s reduced propensity to phase separate. In fact, at much higher RNA concentrations, the LAF-1 RGG C_sat_ can be higher than in the absence of RNA. The concentration at which RNA switches its role as a promoter to a disruptor of protein LLPS may be expected to be near the net neutral state of the solution. However, we see a small shift in this, possibly due to the non-electrostatic, multivalent hydrophobic interactions between nucleotides and amino acids. Previous experimental studies by the Banerjee group (59) have provided remarkable insights into this type of non-monotonic change in C_sat_ with increasing RNA concentration. They showed that in the cases of the FUS/RNA, protamine/RNA mixtures, the system displays a reentrant phase separation behavior, similar to what is observed in the case of LAF-1 RGG. This reentrant behavior emerges from a competition between the attractive and repulsive electrostatic interactions between unlike and like charges, respectively, and depends on the net system charge. Banerjee and co-workers also developed a theoretical approach to explain the observed behavior in terms of a charge inversion mechanism which suggests increasing instability of the condensate as more RNA is added to a mixture beyond a certain point due to charge over screening (19). Their theory predicts a switch in the surface charge of the condensates after the critical point which was confirmed with electrophoretic mobility measurements in experiments.

### Effect of protein sequence charge patterning on RNA partitioning inside the condensed phase

In the last section, we showed how presence of RNA could modulate the LLPS of the LAF-1 RGG sequence. As our CG modelling strategy can be used to study sequence-dependent effects, our next goal was to investigate the role of the protein sequence in modulating RNA-protein LLPS. In our recent work on determining the sequence determinants of LAF-1 RGG phase separation, we showed that the patterning of the charged amino acids can play a critical role in controlling the LLPS properties. Based on earlier work highlighting the role of charged amino acid patterning on the conformational properties of IDPs (60), Sawle and Ghosh proposed a sequence charge decoration (SCD) parameter (61) to quantify the distribution of charged amino acids in a protein sequence. Highly negative SCD values for a protein sequence indicate segregation of positive and negative charges within the sequence and can be observed in sequences having patches of similarly charged amino acids. This generally enhances electrostatic intra and inter-molecular interactions because of the cooperativity of interactions between oppositely charged patches (31). This will result in an enhanced phase separation propensity for sequences with lower SCD values, which was tested successfully by a combination of *in silico, in vitro*, and *in vivo* techniques for the LAF-1 RGG sequence in our recent work (54). Smaller negative, or positive values of SCD indicate a sequence with evenly distributed positive and negative charges, or a large net charge, which would be the case for an unstructured RNA chain.

To study the effects of SCD on a binary mixture of protein and RNA molecules, we use a variant of LAF-1 RGG from our previous work (37) where we had shuffled the sequence and selected an instance where positive and negative charges were highly segregated, having a large negative SCD (Fig. 5A). In the case of shufB, we observe significant accumulation of RNA molecules at the interface between the dense and dilute phases as opposed to a relatively well-mixed WT/RNA condensate (Fig. 5B, C). We then conducted additional simulations in a cubic simulation box of a spherical condensed phase in order to ascertain whether the observed microstructural features may be an artefact of the simulation geometry (Fig. 6, insets). We calculate the protein and RNA radial distribution functions (RDFs) with respect to the center of mass of the condensed phase as a spherical density profile (Fig. 6). As we found previously, the WT LAF-1 RGG condensate formed a well-mixed protein-RNA droplet (Fig. 6A), and the shufB condensate formed a core-shell structure, where RNA mostly forms a "shell” around the “core” consisting of shufB protein (Fig. 6B). This is reminiscent of other examples where sub compartmentalization has been observed (62–64), including ones that show similarly structured organization of different components in a mixture to create sub compartments within a condensate (57, 58).

**Figure 5:**
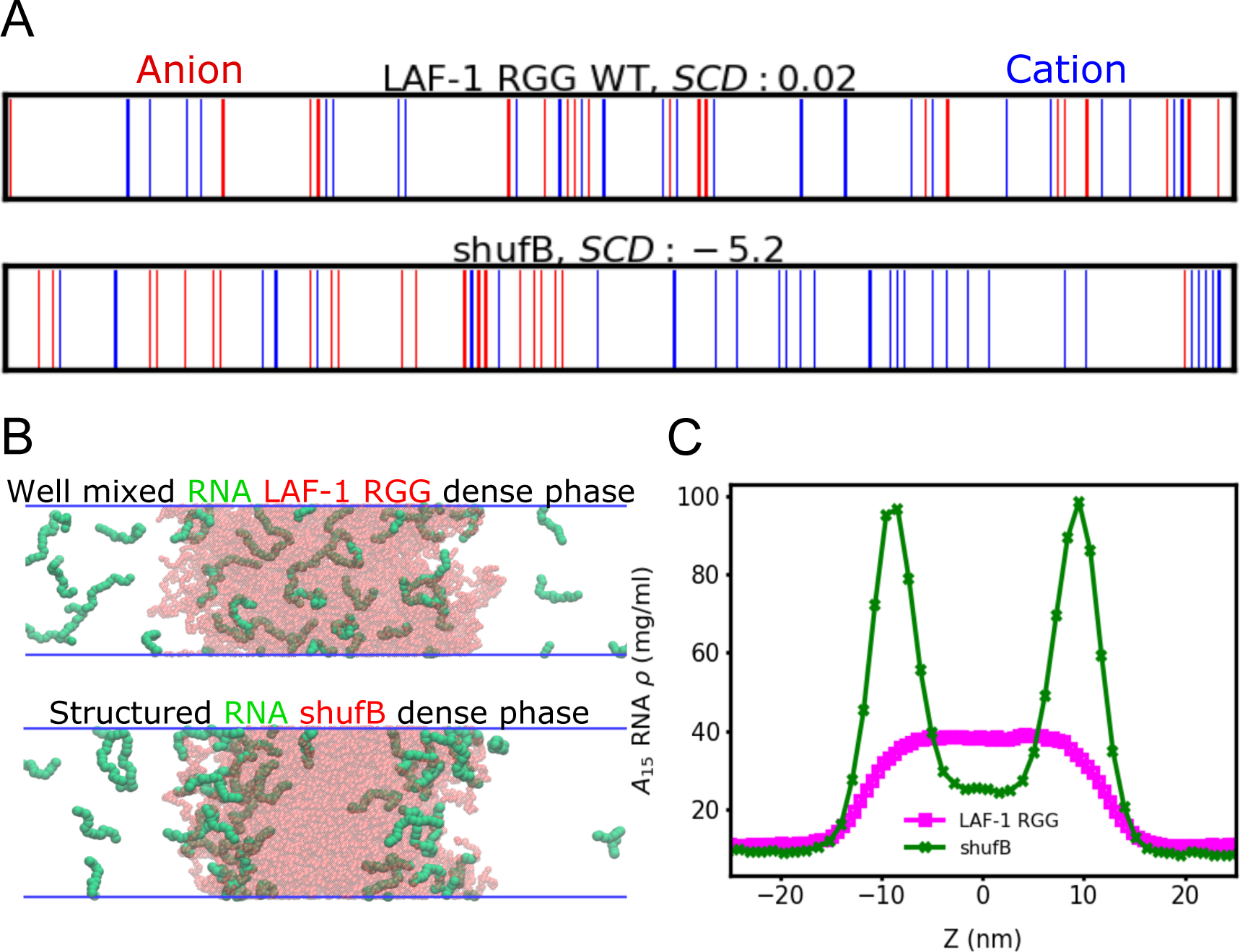
Shuffling protein sequence shows sequence dependence of RNA-protein co-phase separation: Increased charge patterning (lowering SCD) on LAF-1 RGG WT sequence to create shufB (B) Slab snapshots show RNA with preferential localization to the interface for shufB in contrast to the well mixed RNA/LAF-1 RGG case (C) Density profiles from MD simulations at the same simulation temperature and bulk concentrations show sharp interfacial peaks for RNA in shufB whereas RNA in LAF-1 RGG has a central flat peak.

**Figure 6:**
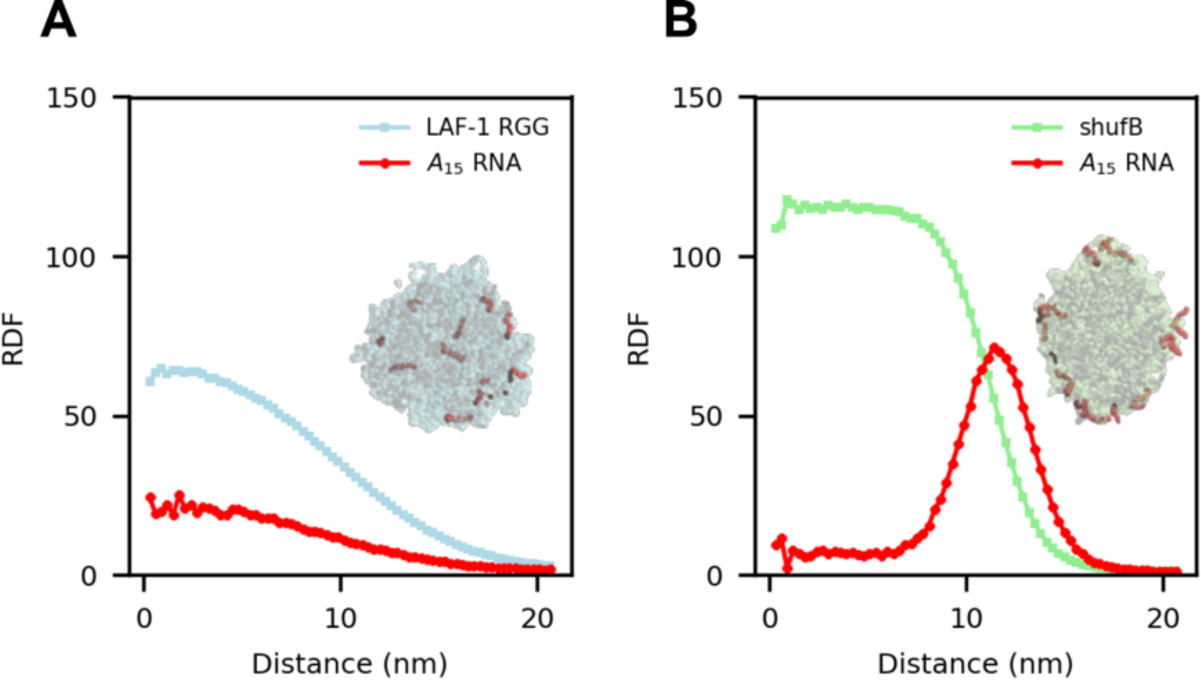
Protein sequence dependent co-phase separation persists in a droplet simulation geometry with intermolecular interactions causing sequence dependent behavior: (A) Radial distribution functions (RDF) from droplet simulations also show a well-mixed condensate for the (A) RNA/LAF-1 RGG mixture and interfacial peaks for the (B) RNA/shufB mixture. Inset images show cross sections of these droplets where we can see RNA molecules (red) located throughout and at the interface for LAF-1 RGG and shufB respectively.

We then ask the question about the driving forces of these two distinct condensate architectures. Our hypothesis is that the observed core-shell architecture emerges from a complex interplay of homotypic protein-protein and heterotypic A_15_ RNA - protein interactions. Looking into the RNA-protein interactions based on the proximity of beads during the simulation we see that the positively charged Arginine residues seem to be making the most contacts (Fig. 7 A) with the negatively charged nucleotides and this is even more pronounced with the shufB sequence as was expected due to the presence of higher charge patterning. Since RNA-RNA interactions will always be repulsive within our model, they would not be expected to contribute to stabilizing of any condensed phase.

**Figure 7:**
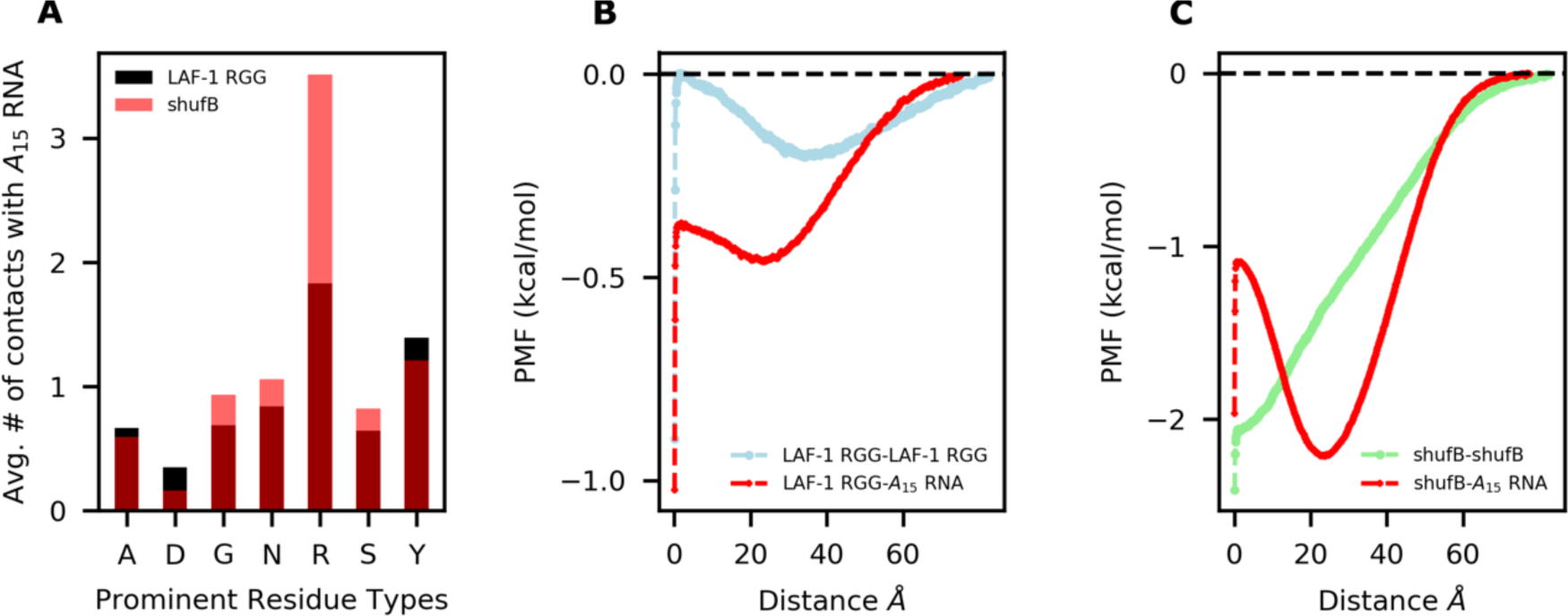
Intermolecular homotypic and heterotypic interactions play a role in modulating RNA-protein co-phase separation: (A) Intermolecular contacts calculated on the basis of proximity of beads in the MD simulation show electrostatic interactions play a major role (B) Potentials of Mean Force for homotypic LAF-1 RGG and heterotypic LAF-1 RGG - RNA interactions and (C) for homotypic shufB and heterotypic shufB - RNA interactions show the formation of a second minima away from zero center of mass distance for shufB - A_15_ RNA interactions linking intermolecular interactions to the observed structure in the shufB-RNA condensate.

We also quantify interactions between a pair of protein-protein or RNA-protein molecules using umbrella sampling Monte Carlo simulations similar to our previous work (30). We thus calculated the potential of mean force (PMF) between the two molecules (protein-protein and A_15_ RNA-protein) as a function of the distance between their centers of mass (Fig. 7 B, C). In all four cases, we see net attractive interactions between the two components, with a strong minimum at zero distance, which is accessible due to the disordered nature of the components. The free energy minimum is deeper in the case of shufB-shufB compared to the LAF-1 RGG – LAF-1 RGG, as would be expected based on their single-component phase behavior (30). We note that most of the PMFs have a secondary minimum at ∼20-30Å, with the exception of shufB-shufB. Interestingly, this second minimum for A_15_ RNA-shufB (∼25 Å) is deeper than the first minimum at zero distance (Fig. 7 C), which is not the case for the other pairs A_15_ RNA - LAF-1 RGG. The homotypic interactions between LAF-1 RGG is similar to the heterotypic interactions between LAF-1 RGG and A_15_ RNA, so A_15_ RNA is well mixed with LAF-1 RGG. However, the homotypic interactions between shufB is quite different from heterotypic interactions between shufB and A_15_ RNA. This is mainly driven by shuffling the LAF-1 RGG sequence towards higher charge patterning, introducing prominent attractive shufB-shufB interactions. This explains the observation that shufB does not form a well-mixed condensate with A_15_ RNA, and that in this case, RNA is majorly situated on the periphery of the shufB condensate (Fig. 5B, C). Comparing how similar the interactions between the homotypic scaffold interactions and the heterotypic scaffold-client interactions might be a useful strategy for checking the microscopic organization in MLOs (65).

## CONCLUSION

In this work we have developed a novel coarse-grained (CG) simulation model for capturing RNA-protein interactions and co-phase separation. We show how this simple RNA model is sufficient to reveal insights into the role RNA plays in modulating protein phase separation. We demonstrate this using the N-terminal disordered RGG domain of the LAF-1 protein, which has been shown to phase separate itself, and in the presence of disordered RNA molecules (23). Our simulations allow us to quantify the degree to which RNA is incorporated into the condensed protein phase and also show effects of RNA on the phase separation propensity of the protein in concordance with experiments (19).

In addition, as our protein model captures the sequence level detail of the disordered protein, it allows us to identify the role that the protein sequence plays in modulating RNA-protein co-phase separation. While keeping the overall amino acid composition constant we shuffled the LAF-1 sequence to identify variants with enhanced electrostatic interactions by creating a more charge-segregated protein sequence. We find that the enhanced electrostatic interactions resulting from this shuffling causes the protein to phase separate more strongly. Interestingly, even though protein-RNA interactions are also enhanced by this charge segregation, we do not observe an increase in RNA incorporation into the condensed phase. Instead, we see interfacial peaks indicating RNA accumulating around the shuffled protein condensate forming a layer between the molecule rich and molecule depleted vapor region in our simulations, and a lower overall concentration of RNA in the condensed phase.

We then looked at 2-molecule interactions by calculating the PMFs between the different molecular pairs. We propose that a complex interplay of homotypic and heterotypic interactions controls co-phase separation leading to the formation of structured droplets. The different structuring of the condensates can be explained by the difference in heterotypic shufB-RNA interactions, and homotypic shufB-shufB interactions, which increase to a greater degree upon shuffling, while heterotypic and homotypic interaction strengths are more balanced in the case of unshuffled LAF-1. Hence, our work here suggests the possibility of using charge patterning of protein sequence to control not only the phase diagram of co-phase separation of proteins and RNA molecules, but also introduce nonhomogeneous microstructures within the liquid droplet.

Even though our simplistic unstructured CG representation of RNA provides us with results in significant agreement with experiments, there are still some effects which one might not be able to capture due to the complex nature of interaction modes which can allow RNA to interact with proteins and itself. Hence, further work on this RNA CG representation is warranted as we would like to capture the more nuanced RNA interactions arising from the π character and secondary structure of RNA molecules which would also play important roles in modulating RNA-protein co-phase separation. Other aspects like improving the charge representation in the CG model can also be important as we show above that electrostatic interactions are a crucial part of how RNA interacts with proteins.

